# Random forest classifiers trained on simulated data enable accurate short read-based genotyping of structural variants in the alpha globin region at Chr16p13.3

**DOI:** 10.1101/2023.11.27.568683

**Authors:** Nancy F. Hansen, Xunde Wang, Mickias B. Tegegn, Zhi Liu, Mateus H. Gouveia, Gracelyn Hill, Jennifer C. Lin, Temiloluwa Okulosubo, Daniel Shriner, Swee Lay Thein, James C. Mullikin

## Abstract

In regions where reads don’t align well to a reference, it is generally difficult to characterize structural variation using short read sequencing. Here, we utilize machine learning classifiers and short sequence reads to genotype structural variants in the alpha globin locus on chromosome 16, a medically-relevant region that is challenging to genotype in individuals. Using models trained only with simulated data, we accurately genotype two hard-to-distinguish deletions in two separate human cohorts. Furthermore, population allele frequencies produced by our methods across a wide set of ancestries agree more closely with previously-determined frequencies than those obtained using currently available genotyping software.

## Background

The *α*-thalassemia genetic trait consists of any of a group of mutations, mainly deletions, that occur within the *α*-globin locus in the human chromosome 16 (16p13.3)(1,2). In general, these mutations result in reduced expression of the *α* globin genes *HBA1* and *HBA2*, and can sometimes lead to severe anemia requiring blood transfusion or even cause death in-utero(3,4). There is evidence that loss of any one of the four copies of the *α* globin genes, resulting in an *α*-thalassemia silent carrier state, is protective against severe malaria, suggesting that higher allele frequencies observed for these mutations in tropical and subtropical climates (malaria belt) may be the result of natural selection(5,6).

The most common *α*-thalassemia deletion is the -*α*^3.7^ deletion. This deletion of approximately 3800 base pairs, and is created by nonhomologous recombination (NHR) between similar sequences known as Z boxes, resulting in the loss of the genomic region between *HBA2* and *HBA1* and the creation of a single hybrid *HBA* gene. Similarly, a less common *α*-thalassemia deletion known as -*α*^4.2^ occurs when NHR of similar sequences called X boxes occurs, resulting in the deletion of about 4200 base pairs including the entirety of *HBA2*. Because the -*α*^4.2^ deletion lies upstream of the -*α*^3.7^ deletion, it is often referred to as the “leftward” deletion, and similarly the -*α*^3.7^ deletion is referred to as the “rightward” deletion(7).

Detection of these and other structural variants (SVs) in the *α*-globin region using short-read whole genome sequence (sr-WGS) data is challenging due to the presence of multiple sequence repeats. These repeats lead to inaccurate alignment of short reads to genomic references such as GRCh38(8) and CHM13v2.0(9) (i.e., aligning reads from the *α*-globin region to the wrong position in the reference), confounding traditional methods for detecting and genotyping SVs.

In recent years, there have been numerous applications of machine learning methods to the characterization of genomes based on short- and long-read DNA sequence data(10–12). These methods have been highly successful at calling variants throughout the genome, but because they are trained on genome-wide data and rely on accurately-aligned reads to make their predictions, they may not perform as well in repetitive regions. In addition, alignment-free methods for genotyping medically relevant variants have been developed(13–15), but so far these have been more applicable to calling smaller variants.

In this work, we present a method that uses simulated sr-WGS data to train random forest (RF) classifiers to call SV genotypes in the *α*-globin region of human chromosome 16. Remarkably, in addition to achieving high accuracy on held-out simulated data, these classifiers are also highly accurate when they are used on sr-WGS from human samples. In two cohorts consisting of 586 and 215 patients, our methods achieve greater than 92% and 99% agreement, respectively, with -*α*^3.7^ genotypes measured by droplet digital PCR (ddPCR). In addition, our method successfully detected three instances of the rare -*α*^4.2^ deletion, indicating that this method could be used as an accurate and valuable screening tool for rare but difficult-to-detect structural mutations. When run on an expanded 1000 Genomes Project (1KGP) cohort of 3,202 samples, our classifier shows high accuracy as measured by the predicted *de novo* mutation rate in 602 trios. Finally, our method’s calls predict allele frequencies across different populations that more accurately mirror known rates of *α*-thalassemia than calls using traditional SV calling methods.

## Results

### Overall accuracy of random forest models when genotyping simulated read sets

In training our random forest model, we used 5-fold cross-validation to optimize our ensemble’s hyperparameters (see “Methods”), then trained the model using these hyperparameters, and finally tested the trained model on held out, simulated samples before using the model on WGS data from real patients (see the following two sections).

On the held out simulated data, trained models easily achieve high accuracy in predicting simulated genotypes. For our random forest classifier, training with 240 samples, each with 30x read coverage, and then testing on held out simulated data with the same coverage resulted in 97.8% mean prediction accuracy (standard error 0.3%). When stratifying by the simulated samples’ depth of coverage, we observe that, as expected, prediction accuracy is significantly lower when samples have lower read coverage (Figure 1A). Models trained on 360 simulated samples with only 10x coverage each have mean prediction accuracy of only 98.3%, while with read coverage of 40x or above, their accuracy is at least 99.5%. WGS datasets with low depth of coverage also present challenges to traditional variant calling software due to a lack of ability to distinguish heterozygous from homozygous genotypes. For example, we observed that the majority of errors for all read coverage values are in samples for which the model confuses the wild type genotype (no deletions on either haplotype) with the heterozygous form of the *α*^4.2^ deletion.

**Figure 1:**
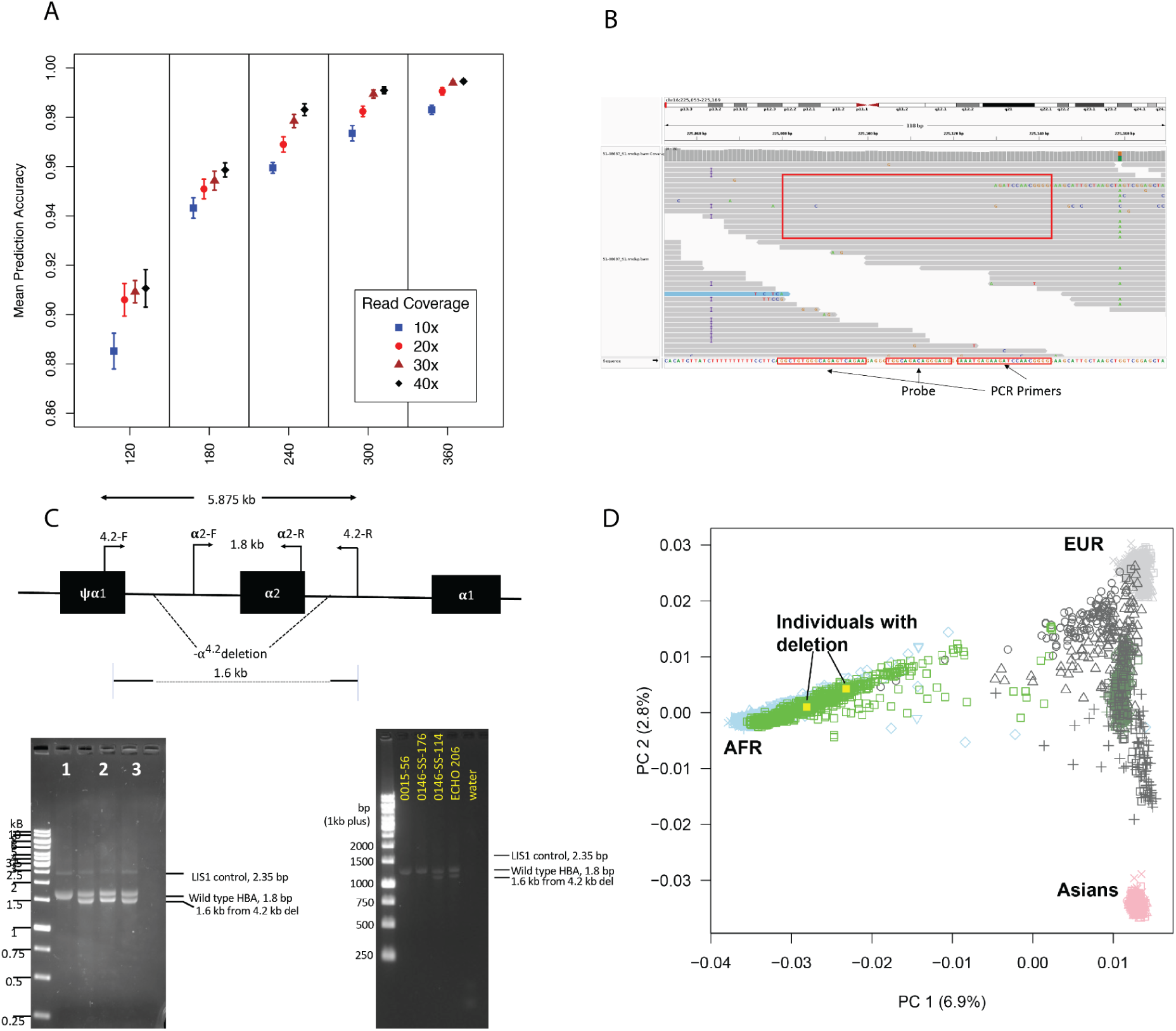
Characteristics of random forest genotyping of alpha thalassemia deletions. A. Plot showing mean accuracy of random forest genotype predictions on held out simulated data vs. number of samples used for training and read depth of coverage per sample. Error bar lengths are standard error values. B. Read coverage in the region of a sample from the first cohort genotyped as homozygous deleted (-*α*^3.7^/-*α*^3.7^) in the ddPCR experiments. Both the amplification primers and the central probe show exactly matching read coverage, indicating possible error in the ddPCR genotyping. C. Gel images of gap PCR confirmation for three out of four samples predicted to be heterozygous for the *α*^4.2^ deletion by the random forest model. D. Principal component analysis of the two individuals in the first cohort confirmed to have the *α*^4.2^ deletion. These two individuals, marked with yellow squares, are shown to be of African ancestry.

### Model performance on WGS data from 586 real individuals

We tested our fitted model by running it on features from a set of 586 FASTQ-formatted files obtained by short read whole genome sequencing of patient samples (cohort 1) which had previously been genotyped by digital droplet PCR (ddPCR) as homozygous wildtype (*αα*/*αα*), heterozygous (*αα*/-*α*^3.7^), or homozygous deleted (-*α*^3.7^/-*α*^3.7^) for the *α*^3.7^ deletion. Even though we allowed the RF classifier to predict any of the six genotypes it was trained to recognize, it classified all but two of the samples as having one of the three *α*^3.7^ genotypes listed above. The remaining two samples were predicted by the model to be heterozygous for the *α*^4.2^ deletion (see “Detection of -*α*^4.2^ in two African American individuals”), but they were genotyped as homozygous wildtype (*αα*/*αα*) in the ddPCR experiment since the *α*^4.2^ deletion was not assessed.

Of the 586 samples, the RF classifier correctly genotyped 539, or 92.0% (Table 1). Of the classifier’s 47 genotyping errors, 30 (63.8%) were in samples for which the predicted genotype was wildtype *αα*/*αα* and the ddPCR showed homozygous deleted (-*α*^3.7^/-*α*^3.7^). Manual inspection of the read data for these samples revealed that nearly all of them contained multiple aligned reads along both (Watson and Crick) strands covering the ddPCR amplicon and probe, with sequence exactly agreeing with the consensus from which the ddPCR primers were designed (Figure 1B). For this reason, we propose that the ddPCR may have failed for these samples. If this is the case, the RF classifier’s accuracy would be 569 correct out of 586, or 97%.

**Table 1:** Comparison of random forest-predicted genotypes to ddPCR genotypes.

|  | Random Forest-Predicted Genotype |  |  |  |
| --- | --- | --- | --- | --- |
| ddPCR Genotypes (First Cohort) | | $-\alpha^{3.7}/-\alpha^{3.7}$ | $\alpha\alpha/-\alpha^{3.7}$ | $\alpha\alpha/\alpha\alpha$ |
| | $-\alpha^{3.7}/-\alpha^{3.7}$ | 8 | 8 | 18 |
| | $\alpha\alpha/-\alpha^{3.7}$ | 1 | 175 | 17 |
| | $\alpha\alpha/\alpha\alpha$ | 0 | 3 | 356 |
| ddPCR Genotypes | | $-\alpha^{3.7}/-\alpha^{3.7}$ | $\alpha\alpha/-\alpha^{3.7}$ | $\alpha\alpha/\alpha\alpha$ |
| (Second Cohort) | $-\alpha^{3.7}/-\alpha^{3.7}$ | **1** | 0 | 0 |
| | $\alpha\alpha/-\alpha^{3.7}$ | 0 | **46** | 0 |
| | $\alpha\alpha/\alpha\alpha$ | 0 | 1 | **66** |

### Model performance on WGS data from a second cohort of 215 real individuals

An additional cohort of 215 individuals (cohort 2) was genotyped using our model to further confirm its accuracy. From this cohort, 114 individuals predicted to have wildtype (68), heterozygous -*α*^3.7^ deletion (47) or homozygous -*α*^3.7^ deletion (1) genotypes were selected for genotyping by ddPCR. Two of the individuals predicted to have the wildtype genotype were found to have an amplification (the “anti-3.7” duplication) and so were not included in our accuracy calculation. Of 112 samples, 111 (99.1%) showed concordance between the ddPCR-determined genotype and the genotype predicted by our RF classifier. Results are presented in Table 1.

Table 1. Confusion matrix of counts comparing genotypes predicted by the random forest model and genotypes determined by digital droplet PCR for the first and second cohorts of samples. For the second cohort, two samples determined by ddPCR to have the anti-3.7 duplication were not included in the counts. Bolded counts represent the counts of correct predictions.

### Detection of -*α*^4.2^ in three African American individuals

Two African American individuals in the first cohort of 586, as well as two additional samples in the 2nd cohort of 215, were predicted by the random forest model to be heterozygous for the -*α*^4.2^ deletion. The -*α*^4.2^ deletion is rare in African populations(7,16), so the existence of samples which are predicted to have the mutation was unexpected. Gap PCR experiments (Figure 1C) confirmed that the machine learning model correctly predicted the -*α*^4.2^ deletion in three of the four individuals. As expected, ancestry analysis of the three samples that were confirmed to have the deletion indicates that they cluster midway between African and European 1000 Genomes Project samples (Figure 1D). Although the fourth individual predicted by our model to have the -*α*^4.2^ deletion was likely a false positive, we investigated the possibility that this individual harbors a larger deletion that was not included in the training set for our model.

### Comparison of random forest-predicted genotypes to those of traditional SV genotyping software

To investigate how our machine learning-based approach to SV genotyping compares to current best practices in short read SV detection and genotyping, we ran our random forest model on WGS short-read data for 3,202 1KGP samples(17). In this dataset, each sample was sequenced to high coverage (targeted 30x). The entire set of 3,202 samples included 2,504 unrelated individuals, with an additional 698 related samples that resulted in the inclusion of 602 family trios. Byrska-Bishop et al. distributed structural variant calls in VCF format for all 3,202 samples, which they obtained by integrating calls from three algorithms: GATK-SV, svtools, and Paragraph (18–20).

While the Byrska-Bishop call set predicts both the -*α*^3.7^ and the -*α*^4.2^ deletions, it detects these two deletions in only a very small minority of the 3,202 individuals, reporting 31 heterozygotes and one homozygous sample for the -*α*^3.7^ deletion, and just six heterozygotes for the -*α*^4.2^ deletion. This corresponds to an allele frequency for the -*α*^3.7^ deletion of less than 1% in all five of the “superpopulations” sequenced for that study, and to far lower frequencies for the -*α*^4.2^ deletion.

However, the frequency of the -*α*^3.7^ deletion, in particular, is known to be quite high in tropical and subtropical regions(21). While the -*α*^3.7^ deletion is less common in European populations, previous studies have estimated its frequency at 8.5% in northeastern Malaysia(22) and 24.8% in Afro-Brazilians(23), so the genotyping methods of Byrska-Bishop et al. are almost certainly characterized by a high false negative rate when detecting these variants.

When run on the 3,202 samples’ WGS data, our random forest classifier predicted that 443 of the samples carried one or two copies of the -*α*^3.7^ deletion (402 heterozygotes and 41 homozygotes), a number far higher than the 32 individuals predicted by Byrska-Bishop et al. Our model also predicted heterozygous -*α*^4.2^ deletions in 32 individuals, compared to just six heterozygotes in the calls of Byrska-Bishop et al. By examining only calls for the 2,504 unrelated samples, we were able to estimate the allele frequency of the two deletions in each of the five superpopulations, obtaining estimates for -*α*^3.7^ ranging from less than 1% in Europeans to greater than 20% in Africans. Estimates for the -*α*^4.2^ population frequency ranged from 0.1% in Europeans to nearly 2% in East Asians (Table 2).

**Table 2:**
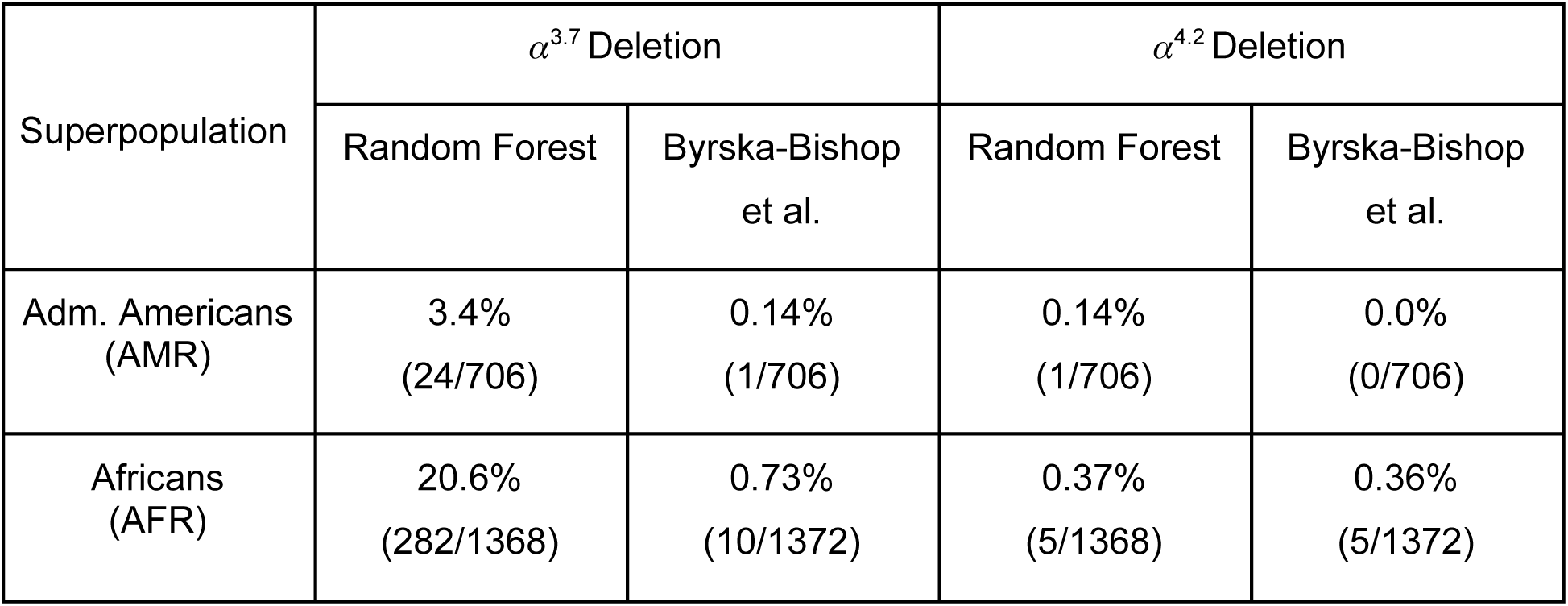
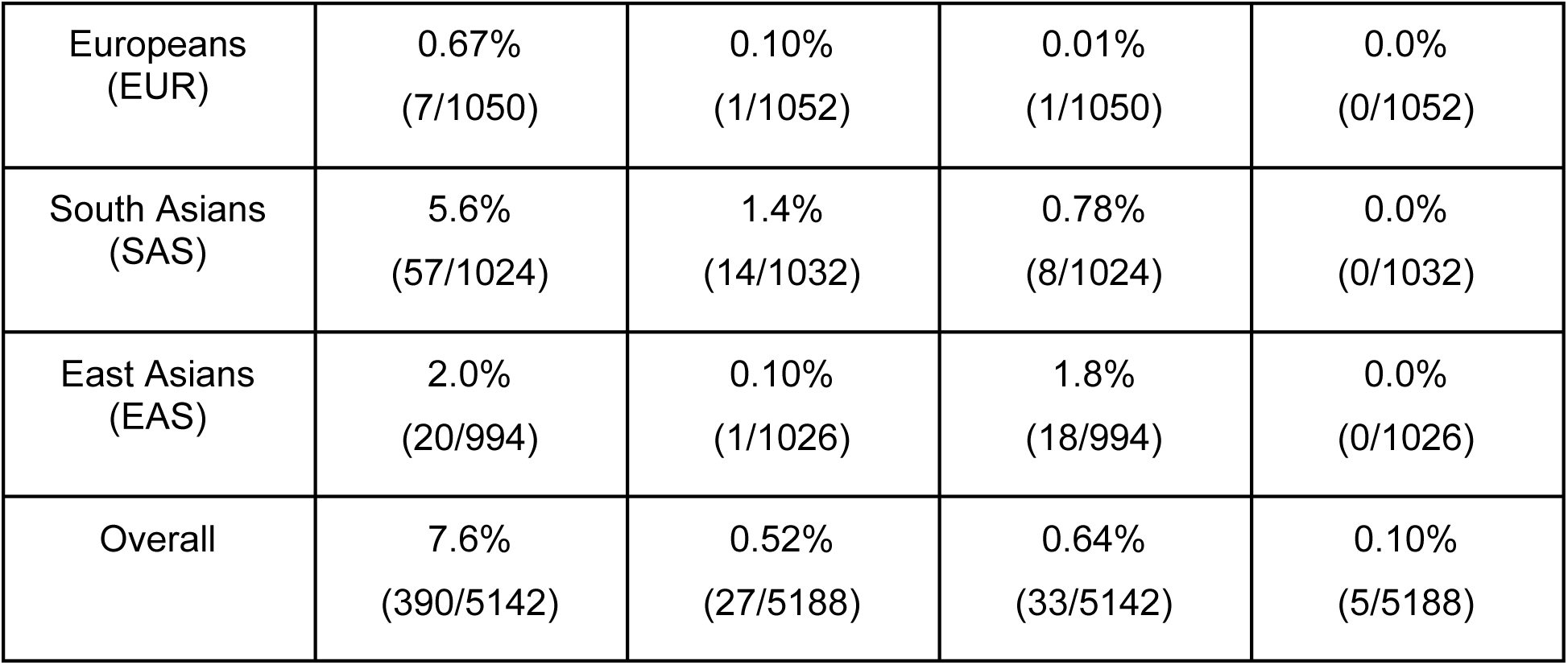
Predicted allele frequencies for alpha thalassemia deletions in the thousand genome superpopulations.

To assess the accuracy of our calls, we examined genotypes predicted for the children of the 602 included family trios, and determined whether their two predicted alleles adhered to Mendelian inheritance rules when compared to those of their two parents. In other words, we confirmed that one of the child’s two predicted alleles was also predicted in one parent, and the other of the child’s two alleles was predicted in the other parent. Only one child out of the 602 (approximately 0.2%) had a predicted genotype that showed Mendelian inconsistency. This sample was of East Asian descent, and had a predicted genotype of -*α*^3.7^/-*α*^4.2^, which we believe might actually be one of the larger deletions present in Asia which our ML models are not yet trained to recognize (see Discussion).

Table 2. Predicted allele frequencies of the -*α*^3.7^ and the -*α*^4.2^ deletions for the 2,504 unrelated 1000-Genomes individuals sequenced to 30x coverage and genotyped for structural variation in Byrska-Bishop et al.

## Discussion

The results we report here show the feasibility of characterizing difficult regions of the human genome using hard-to-align short sequencing reads by training machine learning classifiers to recognize signatures of known variants. Importantly, we have shown that we are able to predict genotypes successfully in *real* sample data using models trained only with *simulated* read data, suggesting that simulated short read data captures features which correlate well with different genotypes.

While in the future the characterization of human genomes will almost certainly be by comparison to an entire pangenomic reference(24), many large short read WGS datasets already exist and are publicly available, without available tools to interrogate investigators’ regions of interest. The work here offers methods for scanning these large datasets for variants effectively without using less accurate, traditional methods for short aligned reads, or conducting expensive experiments.

The *α*-globin region is the site of numerous other, larger deletions which, although they are more rare than the two that we studied, are also medically relevant. While we felt it was beyond the scope of this study to include them in our work, our methods are likely to be successful in distinguishing these different variants.

For regions other than the *α*-globin region, our publicly available software packages, mlfeatures and mlgenotype, allows users to train and use random forest classifiers to recognize SVs across the genome. Interrogating different genomic regions will allow us to determine whether these types of machine learning methods present a viable alternative to traditional structural variant calling methods for using short reads to infer SV genotypes in difficult regions of the human genome, whether it be from a single-haplotype or a pangenomic reference.

### Conclusions

The machine learning method presented here advances state-of-the-art analysis of genomic structural variation using short read sequencing data. Unlike methods based on aligned reads, this method uses the sequence from the reads themselves to classify sequenced samples into known genotypes, and unlike most supervised machine learning methods, it produces accurate results when trained using only simulated data. In addition, these methods can be adapted to classify samples using a pangenomic reference, allowing reanalysis of numerous existing large data sets.

## Methods

### Training classifiers to recognize region-specific read mapping patterns for SV genotypes

In genomic regions such as the *α*-globin region on human chromosome 16 (16p13.3), where the reference contains repeats comparable in length to the length of short sequence reads, mapping of reads to a reference is generally inaccurate, resulting in misplaced reads and multi mapped reads with low mapping quality scores. As a result, software tools which use these aligned reads to predict and genotype SVs will exhibit lower accuracy in these regions, because the aligned reads won’t exhibit the expected patterns that large insertions and deletions generally create in a sample’s data.

However, even if read mappers are inaccurate for samples with SVs, the locations to which reads are mapped in these samples are in most cases consistent, so that different SV genotypes can reasonably be expected to produce predictable patterns of sequence coverage, even if these patterns aren’t easily recognizable by standard SV calling software or manual inspection. Motivated by the hypothesis that these genotype-associated patterns exist, we trained machine learning classifiers by generating simulated sequence reads from a diverse set of diploid genome references mutated to contain the two most common *α*-thalassemia deletions on zero, one, or both copies of *HBA1* and *HBA2* on chromosome 16, and then used these reads’ k-mer compositions to train random forest classifiers to recognize the different genotypes.

Using features (in this case, k-mer counts) calculated from the simulated read data for six possible genotypes (*αα*/*αα*, *αα*/-*α*^3.7^, *αα*/-*α*^4.2^, -*α*^3.7^/-*α*^3.7^, -*α*^4.2^/-*α*^4.2^, and -*α*^3.7^/-*α*^3.7^), we trained random forest classifiers to predict those genotypes. We then tested our model on held out read data from simulated datasets with known deletion genotypes, as well as on sr-WGS datasets from 2 cohorts of African descent; cohort 1 was enrolled under protocols 01H088 /NCT00011648 and 04H0161/ NCT00081523 and comprised patients with sickle cell disease, while cohort 2 was enrolled under protocol 18-H-0146/NCT03685721 and included subjects with sickle cell disease, healthy controls and sickle cell carriers. We also ran our genotype prediction model on the 2,504 samples from the expanded 1000 Genomes Project.

### Simulation of training data from a diverse set of diploid genomes

To train machine learning models to recognize known deletions and amplifications in the *α*-globin region, we first used the ART simulator(25) to simulate FASTQ-formatted reads with specified read length, insert size distribution, and sequence error profiles. For each simulated read set, we used as our genome consensus two constructed haplotypes: one maternal and one paternal haploid region taken from seven diploid assemblies (14 haplotypes) assembled by the Human Pangenome Reference Project(24). For each of these 49 possible haplotype combinations, we introduced -*α*^3.7^ and -*α*^4.2^ mutations to one, both, or neither of the haplotypes to create diploid FASTA files for each of the six possible homozygous or heterozygous genotypes. We then ran custom-built read simulation software on our high performance computing cluster (HPCC), using a Snakemake pipeline, available at https://github.com/nhansen/mlgenofeatures.

### K-mer count features used for training

A “k-mer” is a k-length subsequence of a genomic region. To use k-mers as features for training a random classifier, we computed k-mer count distributions from our read datasets using the k-mer database tool “meryl”(26), generating feature sets for each of our simulated read sets from 49 diverse individuals for each genotype we trained our model to recognize (49 haplotype combinations x 6 genotypes x number of replicates). Because of the large number of k-mers represented in our region of interest (over 40,000 for the *α*-globin region) and the lack of independence of k-mer counts for overlapping k-mers, we subsampled the k-mer counts, using only the counts from every 30th k-mer as features.

### Training neural network and random forest classifiers

To train our random forests (RF) as classifiers to predict genotypes using k-mer counts from simulated and aligned read data, we implemented a python script using the RandomForestClassifer class of the scikit-learn library(27). For all models, training and hyperparameter fitting were performed using 5-fold cross validation on purely simulated data, and the cross validation accuracy estimates were checked against 100 held out simulated samples. Once each classifier was completely fitted using the simulated data, it was used to predict genotypes using features generated from either simulated reads or WGS BAM files from real patients. The python code used for creating simulated datasets is available at https://github.com/nhansen/mlgenofeatures, and code for training and testing all models in this manuscript is available at https://github.com/nhansen/mlgenotype. Further details and a description of how to use the software to genotype other genomic regions is included in the Supplementary Methods.

### Droplet digital PCR genotyping of the 3.7 deletion

Digital PCR for detection of 3.7 kb HBA deletion was carried out using Bio-Rad droplet digital PCR system (QX200 Droplet Digital PCR System, Bio-Rad Laboratories, Hercules, CA). The reference gene was RPP30 amplicon, probe labeled with HEX (part #: 10031244, HEXUniqueAssayID: dHsaCP2500350, Bio-Rad Laboratories). The primers for HBA 3.7 kB deletion detection are forward primer: 5’-GGCTGTGGGCAGAGTCAGAA-3’, reverse primer: 5’-CCCCGTTGGATCTTCTCATTT-3’, the probe is 5’-TGGCAGACAGGGAGG-3’ labeled with FAM and quenched with Iowa black. The reaction mixture consisted of 10 microliters of 2X ddPCR Supermix for Probes (No dUTP), one microliter of 20x primers/probe (900 nM primers, 250 nM probes) and one microliter of 20x RPP30 primers/probe in a total reaction volume of 20 microliters using 20 ng of genomic DNA. The droplets were generated with Automated Droplet Generator (Bio-Rad laboratories). The PCR mixture was then subjected to a cycling condition with an initial denaturation step of 95C for 10 min, followed by a total of 40 cycles, each of which consists of 30 sec denaturation step at 94C, annealing/extension temperature at 60C for 1 min, and at the end of PCR a final denaturation step at 98C for 10 min. After the thermal cycling process, the sealed 96-well plate was placed in the QX200 Droplet Reader for data acquisition and analysis. The copy number of alpha hemoglobin gene was determined by relative to the RPP30.

### Gap PCR for Detection of the 4.2 kb deletion on HBA complex

Gap PCR was performed using Qiagen multiplex PCR reagent (Qiagen, ID: 206143), according to the procedure reported^1, 2^. The design of the reaction (Figure 1A) includes sites of primers for normal (α2-F and α2-R) with PCR product of 1.8 kb and flanking the 4.2 kb deletion (4.2-F and 4.2-R) giving rise to a PCR product of 1.6 kb when the deletion is present. Control for PCR reaction is the reference PCR gene, LIS1, with PCR product of 2.35 kb. Primers 4.2-F and 4.2-R produces a PCR product of 5.875 kb on wild type *HBA* that is too large for PCR cycling conditions. However, the product will be 1.6 kb when the deletion is present. The heterozygous 4.2 kb deletion of *HBA* control DNA was kindly provided by Dr. Matthew Oakley from Viapath, King’s College Hospital, UK.

The primers for control gene LIS1 are, LIS1-F, 5’-ATACCATGGTTACCCCATTGAGC-3’ and, LIS1-R, 5’-AGGGCTATTACATGTGGACCC-3’. The primers for wild type alpha hemoglobin gene are, α2-F, 5’-CCCCTCGCCAAGTCCACCC-3’ and, α2-R, 5’-AGACCAGGAAGGGCCGGTG-3’. The gap PCR primers for 4.2 kb deletion are 4.2-F, 5’-GGTTTACCCATGTGGTGCCTC-3’ and 4.2-R, 5’-CCCGTTGGATCTTCTCATTTCCC-3’. One hundred microgram genomic DNA was included in the reaction. The PCR reaction mixture was incubated at 95°C for 10 min, followed by 35 cycles of PCR, each cycle consisting of: 97°C for 45 sec, 60°C for 1 min 15 sec, and 72°C for 2 min 30 sec. At the end of reaction, the reaction was incubated at 72°C for 5 min. Fifteen microliters of the reaction mix were analyzed in 2% agarose gel.

### Ancestry inference for 4.2 deletion carriers

First, we performed quality control analysis within the cohort of African Americans, filtering by minor allele frequency (--maf 0.01), per genotype missingness (--geno 0.05), and deviation from Hardy Weinberg equilibrium (--hwe 1×10-8). We also pruned strand-ambiguous SNPs and SNPs in high linkage disequilibrium (--indep-pairwise 50 10 0.8). Second, the African american cohort data was merged with the 1000 Genomes Project Phase 3 (PMID: 26432245), and we performed principal component analysis (PCA) using EIGENSTRAT (PMID: 16862161).

## Abbreviations

NHR: non-homologous recombination
SV: structural variant
sr-WGS: short-read whole genome sequence
ddPCR: droplet digital PCR
RF: random forest
1KGP: 1000 Genomes Project

## Declarations

## Ethics approval and consent to participate

“Cohort 1” was enrolled under protocols 01H088 /NCT00011648 and 04H0161/ NCT00081523 and comprised patients with sickle cell disease, while “Cohort 2” was enrolled under protocol 18-H-0146/NCT03685721 and included subjects with sickle cell disease, healthy controls and sickle cell carriers.

## Consent for publication

Not applicable.

## Availability of data and materials

Assemblies used to create templates for read simulation are available from the Human Pangenome Reference Project’s website at the link https://github.com/human-pangenomics/hpgp-data. Instructions for downloading the particular haplotype assemblies used in this study are included in the Supplementary Methods. No other data are needed to simulate reads and train random forest models using the software at https://github.com/nhansen/mlgenofeatures and https://github.com/nhansen/mlgenotype.

The structural variant calls from Byrska-Bishop et al. have been archived by the International Genome Sample Resource (IGSR) and can be downloaded from the link http://ftp.1000genomes.ebi.ac.uk/vol1/ftp/data_collections/1000G_2504_high_coverage/working/20210124.SV_Illumina_Integration/.

## Authors’ contributions

NFH wrote the manuscript. NFH, ZL, MHG, DS conceived of the project and performed analyses. NFH, DS, SLT, and JCM supervised the project. NFH, ZL, GH, and JCL wrote software and evaluated the machine learning models. XW, MBG, and TO ran dd-PCR and Gap-PCR experiments to confirm genotypes. All authors read and approved the final manuscript.

## Acknowledgments

This work utilized the computational resources of the NIH HPC Biowulf cluster (http://hpc.nih.gov). NFH acknowledges Dmitry Antipov for helpful conversation and suggestions regarding this work.

## References

1. Higgs DR. The Molecular Basis of -Thalassemia. Cold Spring Harb Perspect Med. 2013 Jan 1;3(1):a011718–a011718.

2. Farashi S, Harteveld CL. Molecular basis of α-thalassemia. Blood Cells Mol Dis. 2018 May;70:43–53.

3. Harteveld CL, Higgs DR. α-thalassaemia. Orphanet J Rare Dis. 2010 Dec;5(1):13.

4. Gilad O, Steinberg-Shemer O, Dgany O, Krasnov T, Noy-Lotan S, Tamary H, et al. Alpha-Thalassemia Carrier due to –α^3.7^ Deletion: Not So Silent. Acta Haematol. 2020;143(5):432–7.

5. Flint J, Hill AVS, Bowden DK, Oppenheimer SJ, Sill PR, Serjeantson SW, et al. High frequencies of α-thalassaemia are the result of natural selection by malaria. Nature. 1986 Jun;321(6072):744–50.

6. Mockenhaupt FP, Ehrhardt S, Gellert S, Otchwemah RN, Dietz E, Anemana SD, et al. α+-thalassemia protects African children from severe malaria. Blood. 2004 Oct 1;104(7):2003–6.

7. Embury SH, Miller JA, Dozy AM, Kan YW, Chan V, Todd D. Two different molecular organizations account for the single alpha-globin gene of the alpha-thalassemia-2 genotype. J Clin Invest. 1980 Dec 1;66(6):1319–25.

8. Schneider VA, Graves-Lindsay T, Howe K, Bouk N, Chen HC, Kitts PA, et al. Evaluation of GRCh38 and de novo haploid genome assemblies demonstrates the enduring quality of the reference assembly. Genome Res. 2017 May;27(5):849–64.

9. Nurk S, Koren S, Rhie A, Rautiainen M, Bzikadze AV, Mikheenko A, et al. The complete sequence of a human genome. Science. 2022 Apr;376(6588):44–53.

10. Poplin R, Chang PC, Alexander D, Schwartz S, Colthurst T, Ku A, et al. A universal SNP and small-indel variant caller using deep neural networks. Nat Biotechnol. 2018 Nov;36(10):983–7.

11. Cai L, Wu Y, Gao J. DeepSV: accurate calling of genomic deletions from high-throughput sequencing data using deep convolutional neural network. BMC Bioinformatics. 2019 Dec;20(1):665.

12. Shafin K, Pesout T, Chang PC, Nattestad M, Kolesnikov A, Goel S, et al. Haplotype-aware variant calling enables high accuracy in nanopore long-reads using deep neural networks [Internet]. Bioinformatics; 2021 Mar [cited 2023 Sep 4]. Available from: http://biorxiv.org/lookup/doi/10.1101/2021.03.04.433952

13. Boudry A, Darmon S, Duployez N, Figeac M, Geffroy S, Bucci M, et al. Frugal alignment-free identification of FLT3-internal tandem duplications with FiLT3r. BMC Bioinformatics. 2022 Oct 28;23(1):448.

14. Grytten I, Dagestad Rand K, Sandve GK. KAGE: fast alignment-free graph-based genotyping of SNPs and short indels. Genome Biol. 2022 Oct 4;23(1):209.

15. Ndila CM, Nyirongo V, Macharia AW, Jeffreys AE, Rowlands K, Hubbart C, et al. Haplotype heterogeneity and low linkage disequilibrium reduce reliable prediction of genotypes for the -α3.7I form of α-thalassaemia using genome-wide microarray data. Wellcome Open Res. 2021 Sep 22;5:287.

16. Steinberg M, Embury S. Alpha-thalassemia in blacks: genetic and clinical aspects and interactions with the sickle hemoglobin gene. Blood. 1986 Nov 1;68(5):985–90.

17. Byrska-Bishop M, Evani US, Zhao X, Basile AO, Abel HJ, Regier AA, et al. High-coverage whole-genome sequencing of the expanded 1000 Genomes Project cohort including 602 trios. Cell. 2022 Sep;185(18):3426–3440.e19.

18. Collins RL, Brand H, Karczewski KJ, Zhao X, Alföldi J, Francioli LC, et al. A structural variation reference for medical and population genetics. Nature. 2020 May 28;581(7809):444–51.

19. Larson DE, Abel HJ, Chiang C, Badve A, Das I, Eldred JM, et al. svtools: population-scale analysis of structural variation. Valencia A, editor. Bioinformatics. 2019 Nov 1;35(22):4782–7.

20. Chen S, Krusche P, Dolzhenko E, Sherman RM, Petrovski R, Schlesinger F, et al. Paragraph: a graph-based structural variant genotyper for short-read sequence data. Genome Biol. 2019 Dec;20(1):291.

21. Piel FB, Weatherall DJ. The α-Thalassemias. Longo DL, editor. N Engl J Med. 2014 Nov 13;371(20):1908–16.

22. Rosnah B, Rosline H, Zaidah AW, Noor Haslina MN, Marini R, Shafini MY, et al. Detection of Common Deletional Alpha-Thalassemia Spectrum by Molecular Technique in Kelantan, Northeastern Malaysia. ISRN Hematol. 2012 Jul 19;2012:1–3.

23. Alcoforado GH de M, Bezerra CM, Lemos TMAM, Oliveira DM de, Kimura EM, Costa FF, et al. Prevalence of α-thalassemia 3.7 kb deletion in the adult population of Rio Grande do Norte, Brazil. Genet Mol Biol. 2012 Jul 26;35(3):594–8.

24. Liao WW, Asri M, Ebler J, Doerr D, Haukness M, Hickey G, et al. A draft human pangenome reference. Nature. 2023 May 11;617(7960):312–24.

25. Huang W, Li L, Myers JR, Marth GT. ART: a next-generation sequencing read simulator. Bioinformatics. 2012 Feb 15;28(4):593–4.

26. Rhie A, Walenz BP, Koren S, Phillippy AM. Merqury: reference-free quality, completeness, and phasing assessment for genome assemblies. Genome Biol. 2020 Dec;21(1):245.

27. Pedregosa F, Varoquaux G, Gramfort A, Michel V, Thirion B, Grisel O, et al. Scikit-learn: Machine Learning in Python. 2012 [cited 2023 Sep 4]; Available from: https://arxiv.org/abs/1201.0490

