## Supplementary Information for "Random forest classifiers trained on simulated data enable accurate short read-based genotyping of structural variants in the alpha globin region at Chr16p13.3"

### Reproducing this manuscript’s results by running its software on publicly available data

*Installing prerequisite software*

To simulate k-mer-based features for training a random forest classifier, first ensure that you have installed all prerequisite software required by the mlgenofeatures and mlgenotype pipelines:

- Python v3.7 or greater
- Pysam python library (https://pysam.readthedocs.io/en/stable/ )
- Snakemake (https://snakemake.readthedocs.io/en/stable/)
- ART version ART-MountRainer-2016-06-05 (https://www.niehs.nih.gov/research/resources/software/biostatistics/art/index.cfm)
- meryl version 1.3 (https://github.com/marbl/meryl)
- Samtools (http://www.htslib.org/)

*Simulating data from the $\alpha$-globin region for various haplotypes on different genomic backgrounds*

First clone the github repository at https://github.com/nhansen/mlgenofeatures:

```
>git clone https://github.com/nhansen/mlgenofeatures
>cd mlgenofeatures
```

To run mlgenofeatures, you’ll need to download seven assemblies (14 haplotypes, which will require about 20gb of disk space) from the HPGP data repository:

```
>cd resources/HPGP/haplotype_assemblies/
>for url in `cat assembled_hpgp_haplotype_urls.txt`; do \
    export file=`echo $url | sed 's:.*/::'`; \
    curl $url --output $file; \
done
```

The download may take a while, but once it's finished, you should have 14 gzipped fasta files in the haplotype_assemblies directory which you can then convert to bgzipped files and index with samtools:

```
>for fasta in `ls HG*fa.gz`; do \
    gunzip -c $fasta | bgzip -c > $fasta.bgzip; \
    mv $fasta.bgzip $fasta; \
    samtools faidx $fasta; \
done
```

Once the HPGP genome fastas are downloaded into the appropriate directory and indexed, you can configure the Snakemake pipeline by editing the "config.yaml" file in the config directory, and create files of training features using the "sh.make_sim" script.

*About the config file for running mlgenofeatures*

Several useful parameters for simulating the six $\alpha$-thalassemia genotypes are in the config file located at config/config.yaml in the mlgenofeatures repository. These parameters can be tuned to the particular srWGS dataset you are analyzing, so the model will be trained with simulated data that looks like your real data. For example, the parameter "readlength" can be set to the the length of the reads, and "insertlength" can be set to the mean total length of your library inserts (from the start of the first read to the start of the second read), while "insertstddev" sets the standard deviation of insert lengths in the simulated dataset.

By default, mlgenofeatures will generate k-mer features from simulated read datasets with a variety of depths of coverage (ranging from 10x to 60x), but if you know specifically what depth of coverage your whole genome read sets have, you can tailor the simulated read coverage to be closer to your dataset's by editing the list following the line "coveragevals:" in the config file. The rest of the parameters in the config file shouldn't be altered unless you are creating a simulated dataset for a different part of the genome or with different background genome fastas.

*Generating feature files to use in training*

With the Snakefile located in the "workflow" subdirectory of the repository, the snakemake command can be used to create feature files for each of the six $\alpha$-thalassemia genotypes:

```
>cd ../..
>for geno in (WTYP_WTYP, WTYP_AL37, WTYP_AL42, AL37_AL37, \
AL37_AL42, AL42_AL42); do
    snakemake —cores=16
features/combined_feature_file.$geno.txt
done
```

These “combined feature” files can be combined into a single file for use in training classifiers by extracting the header from one of them and combining it with headerless versions of each of the six files:

```
>grep ‘genotype’ features/combined_feature_file.AL37_AL37.txt >
combined_features.sixgenos.txt
>for geno in (WTYP_WTYP, WTYP_AL37, WTYP_AL42, AL37_AL37, \
AL37_AL42, AL42_AL42); do
    grep -v ‘genotype’ features/combined_feature_file.$geno.txt
>> combined_features.sixgenos.txt
done
```

Now we can split the combined file of features into equal sized training and test sets:

```
>awk 'NR==1 || (NR-10*int(NR/10)>=0 && NR-10*int(NR/10)<=4)\
{print}' combined_features.sixgenos.txt >\
combined_features.sixgenos.train.txt
>awk 'NR==1 || (NR-10*int(NR/10)>=5 && NR-10*int(NR/10)<=9) \
{print}' combined_features.sixgenos.txt > \
combined_features.sixgenos.test.txt
```

*Training random forest models*

To use the features you generated in the previous step to train random forest genotyping models, install the mlgenotype python library, either from PyPi or from bioconda using anaconda.

To install mlgenotype with Python's pip installer, first create a virtual environment. Then use pip install to install the latest version of mlgenotype:

```
>python3 -m venv mlgeno_env
>python3 -m pip install mlgenotype
```

The mlgenotype package is also hosted on anaconda and available through the bioconda channel:

```
>conda create -n mlgeno -c bioconda -c conda-forge mlgenotype
>conda activate mlgeno
```

The python script used to create Figure 1A of this manuscript uses the mlgenotype python library, and is in the mlgenotype repository here: https://github.com/nhansen/mlgenotype/blob/main/mlgenotype/figure1calcs.py. It can be run by creating directories output/models and output/predictions, then running the script:

```
>figure1calcs --train combined_features.sixgenos.train.txt
--test combined_features.sixgenos.test.txt --outdir output
--prefix figure1a
```

The files created in the output directory include the model itself ("output/models/figure1a.[number of training samples].rf.model") in pickle format, the optimal hyperparameters and its estimated accuracy ("output/models/figure1a.[number of training samples].rf.stats"), and importance measures of the k-mer features ("output/models/figure1a.[number of training samples].rf.importance.txt").

The "output/predictions" directory contains predicted genotypes for each model on the held out "test" samples whose features were passed to the script with the –test option. These output files were used to calculate the data plotted in Figure 1A.

*Generation of feature files from real sequencing data in BAM files*

The genotype predictions made for samples from Cohort 1, Cohort 2, and the IGSR samples from Byrska-Bishop *et al.* were done using models trained using features generated with the mlfeatures Snakemake pipeline (see the section *Simulating data from the* $\alpha$*-globin region for various haplotypes on different genomic backgrounds)*. Since the sr-WGS data for these samples was in bam format, after putting all bam files in a subdirectory named "testsamples" within the simulation directory, we first converted each bam file to FASTQ format using the bedtools "bamtofastq" command:

```
>mkdir testsamples
>for file in `ls testsamples`; do \
     export FQ=`echo $file | sed 's/\.bam/.fastq/'`; \
     bedtools bamtofastq -i $file -fq $FQ; \
done
```

Then, for each created fastq file in the testsamples directory, we generate a features file:

```
>for file in `ls testsamples/*.fastq`; do \
     export FEATURES=`echo $file | sed
     's/\.fastq/.kmercounts.txt/'`; \
     snakemake –nolock –cores=16 $FEATURES; \
done
```

Deletion genotypes for the samples' feature files are calculated using the mlgenotype library's rfmodelpredict script:

```
>for file in `ls testsamples/*.kmercounts.txt`; do \
     rfmodelpredict --modelfile trainedmodel.model \
```

```
      --featurefile $file; \
done
```
